# Differential effects of conspecific and heterospecific density on the development of *Aedes aegypti* and *Aedes albopictus* larvae

**DOI:** 10.1101/383125

**Authors:** Robert S Paton, Katherine Heath, Anthony J Wilson, Michael B Bonsall

## Abstract

1. Between-species competition shapes the distribution and abundance of populations. *Aedes aegypti* and *Ae. albopictus* are vectors of pathogens such as dengue and are known to compete at the larval stage.
2. The outcome of this inter-species competition has been found to be context dependent, with the strength and direction changing with resource availability and type. We were motivated by this uncertainty, and aimed to elucidate the magnitude and mechanism of competition.
3. We manipulated the larval density of mixed and single species cohorts of larvae, measuring the effects on survivorship and development time. Unlike other related studies, we adjusted the feeding regime so that the per-capita resource availability was kept constant across all density treatments, at a level sufficient for successful development. This ensured that each larvae at least had the opportunity to gain the requisite resources for pupation.
4. Our analysis found that *Ae. aegypti* suffered notably less mortality due to intra- and interspecific competition. For both species, intra- and interspecific competition led to the survival of faster developing individuals, with the exception that slower developing *Ae. albopictus* larvae survived when exposed a combination of both high con- and heterospecific densities.
5. These results show that the competition between *Ae. aegypti* and *Ae. albopictus* can still occur even when resources are theoretically adequate for development. This suggests that larvae can alter resource seeking and consumption parameters when exposed to high densities of conspecifics and heterospecifics, leading to contest competition. Evidence for resource-independent mechanisms of competition such as crowding are also found, as is evidence for the importance of demographic stochasticity in population processes.

## Introduction

Studying the processes driving the occurrence and persistence of populations is central to the field of ecology (Morin, 2011). Species with directly overlapping niches will compete for resources and space, just as they do with conspecifics. These interactions between individuals of the same species (intraspecific) and different species (interspecific) are crucial in shaping ecological communities and can help explain the abundance and distribution of populations (Chesson, 2000; Morin, 2011). Theory predicts that species experiencing intraspecifc competition to a greater degree than interspecifc competition can co-occur, whereas the opposite (interspecific > intraspecific) will lead to competitive exclusion (Armstrong and McGehee, 1980). The extent of intra- and interspecific interactions can change across gradients of biotic (e.g. predator abundance) and abiotic (e.g. resource availability, constancy, type) variables, meaning both competitive exclusion and coexistence can occur in heterogeneous environments or across landscapes and regions (Chamberlain, Bronstein, and Rudgers, 2014; Amarasekare, 2003). For two species competing for the same resource, competition can be for the resource directly and/or indirectly through interference. The former is simply the effect one species will have on the other if it depletes an essential common resource. In the second case, one species inhibits the competitor by using mechanical, signalling or chemical interference (Chesson, 2000). The effects of competition manifest as penalties in life history parameters, such as survival, growth and development. Quantifying the direction, magnitude and mechanism of intra- and interspecific competition has been the objective of a plethora of empirical and theoretical studies (see Chesson (2000) and the references therein). Some opt to take a phenomenological approach, describing the “net-outcomes” of competition, while others focus on finding mechanistic explanations. Here, we are interested in a specific instance of inter-species competition; that between *Aedes* mosquitoes.

Mosquito-borne viruses are a significant cause of mortality and morbidity particularly in the developing world (Bhatt et al., 2013). The recent Zika outbreak in Brazil and an estimated 100 million annual dengue infections motivates the need for a concerted effort to curtail disease transmission (Yakob and Walker, 2016; Bhatt et al., 2013). With the notable exception of yellow fever, vaccine efficacy is poor, and availability and coverage continue to be a problem (Bhatt et al., 2013). This has led to a focus on controlling the Aedes vectors of these diseases, with emphasis on releases of sterile, genetically-modified or *Wolbachia-infected* male mosquitoes designed to suppress wild populations. The effectiveness of these strategies is contingent on a robust understanding of the ecology of the principal disease vectors, Aedes *aegypti* and Aedes *albopictus*. Experimental (Hancock et al., 2016) and theoretical (Yakob, Alphey, and Bonsall, 2008) work has highlighted the importance of ecological processes - such as density-dependent competition - in determining the effectiveness of control.

*Aedes aegypti* originated in Africa, but has now become established in Asia and the Americas (Kraemer et al., 2015; Brown et al., 2014). It is a primary vector of many arboviral diseases, such as dengue, chikungunya and Zika (Black et al., 2002; Chouin-Carneiro et al., 2016). Female Ae. *aegypti* bite during the day and are highly anthropophilic (Scott and Takken, 2012). This means that the bed-net based biting-prevention strategies employed against malarial mosquitoes are ineffective against *Ae. aegypti*.

*Aedes albopictus* is a secondary vector of arboviruses originating from south-east Asia (Gratz, 2004). Invasive in much of its range, *Ae. albopictus* is capable of occupying more temperate environments, with a range extending northward to the USA and southern Europe (Kraemer et al., 2015). The range expansion of *Ae. albopictus* was facilitated by international shipping, where its dormant eggs can survive in transit, often in old tyres (Delatte et al., 2009; Simard et al., 2005). *Ae. albopictus* is generally considered to have a preference for biting outdoors, only feeding opportunistically on humans (Paupy et al., 2009; Richards et al., 2006) (though see Ponlawat and Harrington (2005) and Delatte et al. (2010) for examples of *Ae. albopictus* anthropophily). Despite its host preferences, Ae. *albopictus* has still been implicated in several outbreaks, such as in Hawaii, China and Japan (Paupy et al., 2009). Particularly relevant to this study is the finding that female Ae. *albopictus* were more likely to become infected with dengue when they had experienced increased levels of intra- and interspecific larval competition (Alto et al., 2008). This highlights its potential as a disease transmitter, and has led to increasing concern that it could act as a maintenance or primary vector (Gratz, 2004; Paupy et al., 2010).

### Competing vectors

*Aedes aegypti* and Ae. *albopictus* are competent invasive species, both having established persistent populations on new continents (Brady et al., 2014). The two species have increasingly come into contact as *Ae. albopictus* expanded its range throughout the 20^th^ century. These species are known to share larval habitats of ephemeral freshwater pools. This direct niche overlap makes larval pools a forum for inter-species competition.

Historically, *Ae. aegypti* displaced *Ae. albopictus* from urban areas in Asia (Macdonald, 1956), such as in Calcutta (Gilotra, Rozeboom, and Bhattacharya, 1967). Braks et al. (2003) and Simard et al. (2005) document spatially segregated co-occurrence in Brazil and Cameroon respectively. The same segregation is documented on some islands, such as in Hawaii (Winchester and Kapan, 2013) and Reunion (Bagny et al., 2009), though declines (but not extinctions) of *Ae. aegypti* were also noted. More recent introductions of *Ae. albopictus* into *Ae. aegypti* occupied areas has resulted in a rapid displacement of *Ae. aegypti*. In the mid-1980s, the introduction of *Ae. albopictus* in Texas resulted in a rapid displacement of *Ae. aegypti* across the Southern States, with it persisting only in select cities in Southern Florida (O’Meara et al., 1995). Habitat preference studies suggest that *Ae. aegypti* seems better able to occupy urban environments, while *Ae. albopictus* has a preference for more vegetated areas (Rey et al., 2006).

Several studies have attempted to determine which vector is the superior larval-stage competitor (reviewed in Juliano (2009)). Results are mixed, with some early studies finding *Ae. aegypti* to be the superior competitor (Moore and Whitacre, 1972; Moore and Fisher, 1969) and subsequent studies *Ae. albopictus* (Reiskin and Lounibos, 2009; Juliano, Lounibos, and O’Meara, 2004; Braks et al., 2004). Most crucial is that the strength and directions of inter-species competition was context dependent. For instance, competitive outcomes have been found to change in response to different resource types (Barrera, 1996; Murrell and Juliano, 2008), temperatures (Farjana, Tuno, and Higa, 2012) and habitat constancy (Costanzo, Kesavaraju, and Juliano, 2005). Moreover it has also been shown that sympatric and allopatric populations of *Ae. aegypti* and *Ae. albopictus* can suffer from and exert different levels of interspecific competition (Leisnham et al., 2009). Context-dependent variation in the outcome of *Aedes* competition was corroborated by Juliano’s 2010 meta-analysis of larval competition studies. He concluded that *Ae. albopictus* suffered less interspecies competition than *Ae. aegypti* in habitats with high-quality food, but that they were competitively equivalent in resource poor habitats. The fact that *Aedes* mosquitoes compete across heterogeneous, fragmented habitats, further complicate findings. Across a heterogeneous landscape, differences in competitive outcomes could allow for persistence in some areas but not others (Juliano, 2009; Amarasekare, 2003).

### Mechanisms of larval competition

Resource competition has long been considered the primary driver of intra- and interspecific larval competition between *Aedes* mosquitoes (Dye, 1984a). Indeed this mechanism has been represented as density-dependent competition in models of mosquito population dynamics (Dye, 1984b), including those aiming to inform optimal disease interventions (Yakob, Alphey, and Bonsall, 2008; Bonsall et al., 2010). The functions describing intra- and interspecific competition in such models can be parameterised by studies where single and mixed species cohorts are reared in different densities and measure the effects on life history parameters.

However, Heath et al. *(In review*) highlight that many mosquitoes-focused empiricists conflate the effects of larval resource availability with *all* density-dependant processes. That is to say that many studies hold feeding regimes constant across density treatments (e.g. Reiskin and Lounibos (2009)), reducing the per-captia resource availability (same resources, more individuals). The subsequent effects measured on survival/growth rates/fecundity are all then treated as the result of resource-mediated competition, aggregating other mechanism with it. This is important, as there are there are other mechanisms by which competition could occur. For instance, evidence from Moore and Fisher (1969) and Moore and Whitacre (1972) demonstrated that the development times of *Ae. albopictus* larvae in high-density mixed-species cohorts were significantly lengthened by increased densities of *Ae. aegypti*. As the larvae were not resource-limited, the authors attribute this to the production of a chemical compound termed a growth retardant factor (GFR), thought to be produced by *Ae. aegypti* at high larval densities (though this result could not be repeated by Dye (1982)). Alternatively, evidence from Dye (1984a) suggests that differences in development time could be attributed to the volume the larvae were reared in. This could be due to mechanical interference reducing feeding efficiency, or by jostling to reach the surface to breath.

We are motivated by the degree of uncertainty around the outcome of larval competition between *Ae. aegypti* and *Ae. albopictus*, and the mechanism by which this competition occurs. In this study, we manipulated the conspecifc and heterospecific larval densities of *Ae. aegypti* and *Ae. albopictus*, and recorded the effect this had on larval survivorship and development time. Crucially, unlike many Aedes-focused competition studies, we adjusted the feeding regime so that the per-capita resource availability was kept constant across all density treatments. We specifically chose a feeding regime under which each larvae *theoretically* had the necessary resources to successfully develop. This allowed us to better isolate mechanistic aspects of larval competition, and better observe what components of competition were attributed to direct resource-mediated or interference-mediated competition.

## Methods

### Routine colony maintenance

Colony mosquitoes were kept at the Pirbright Institute in Surrey, UK. Rearing rooms were maintained at 25°C ± 1°C with a 16:8 light:dark cycle. Pirbright’s *Ae. aegypti* colony was established from a line at the Liverpool School of Tropical Medicine, which was originally from West Africa (Macdonald, 1962). Eggs were placed in 150ml of reverse-osmosis water and vacuum hatched for at least one hour. First instar larvae were then transferred into bowls filled with approximately 1 litre of tap water, in densities of around 450 - 600 larvae per bowl. Larvae were fed 0.15-0.2g desiccated beef liver powder (NOW Foods, Bloomingdale, IL, USA) every two days, when the water was also changed. Pupated larvae were decanted into small 100ml tubs of water and placed inside 40cm^3^ Bugdorm insect cages (MegaView Science, Taichung, Taiwan). Adults were maintained on room temperature 10% sugar solution on a cotton disk, and females fed defibrinated horse blood (TCS Biosciences, Buckingham, UK) using a Hemotek membrane feeding system (Hemotek Ltd., Great Harwood, Lancashire, UK). Tubs lined with damp filter paper were put in each cage after blood feeding for egg-laying.

The *Ae. albopictus* line was Italian, and founded from the Rimini strain described in Dritsou et al. (2015). The maintenance protocol was identical to the above, with the exception that *Ae. albopictus* eggs were hatched in a medium of water and 0.3% dried active yeast (Youngs Home Brew Ltd., Bilston, West Midlands, UK).

### Experimental protocols

Single and mixed species cohorts of *Ae. albopictus* and *Ae. aegypti* larvae were reared under different density conditions. Seventh and eighth generation *Ae. aegypti* and *Ae. albopictus* were used, and were all from the colonies described above. However, instead of hatching *Ae. albopictus* using yeast as per the routine protocol, they were hatched in the same way as *Ae. aegypti*. This kept both the procedure the same across each species and ensured that the yeast did not supplement the resources available to *Ae. albopictus*. As this method differed from the standard hatching procedure for *Ae. albopictus*, the larvae were left in the vacuum for at least 3 hours to ensure there was a sufficient yield of both species.

Egg papers were placed in reverse osmosis water and vacuum hatched overnight. Hatched first instar larvae were counted into 50ml Falcon conical centrifuge tubes (Corning, New York, USA) tubes filled to 49ml with water (1ml of food solution made this up to 50ml). The number of larvae in each tube corresponded to the density combinations given in table 1. The density scale was chosen to give roughly an even number of larvae per treatment while also covering a biologically relevant range. Density was manipulated by changing the number of individuals per tube while holding the volume of the tube constant (See Figure 1). Tubes for low density treatments therefore contained fewer individuals than for high density treatments, reducing the sample size. The lower-density tubes were therefore repeated so that the overall number of larvae experiencing each density condition was around 50 (the number in the highest density treatment). The densities given in the table 1 were explored for each species in isolation (1:0), equal ratios of each species (1:1) and biased ratios (2:1 and 1:2, 1:3 and 3:1). The procedure described in Figure 1 was repeated twice for each ratio. For low-density, mixed-species treatments some of ratios could not be achieved with number of individuals in the tube (e.g. 4:3 can not be configured from 6 larvae). The density of these treatments was therefore increased slightly in order to allow for these configurations.

**Table 1:** A description of the density conditions (in larvae per millilitre, La/ml) explored in each treatment (T1 - 5). The quoted ratios are those of *Ae. aegypti* to *Ae. albopictus*. Each condition/ratio combination was repeated twice. Tubes are repeated so that there were approximately 50 larvae subjected to each of the density treatments. Food treatments are scaled to the number of larvae in each pot (milligrams per pot, mg/pot) from the amount decided on in the pilot study. In low-density treatments, 6 individuals were insufficient to achieve the required ratios. We therefore increased the number of larvae to the minimum amount needed to make the ratio, and adjusted the feeding regime accordingly.

|  | Density Treatment |  |  |  |  |
| --- | --- | --- | --- | --- | --- |
|  | T1 | T2 | T3 | T4 | T5 |
| **Larvae per tube (La)** | 6 | 16 | 28 | 38 | 50 |
| **Density (La/ml)** | 0.12 | 0.32 | 0.56 | 0.76 | 1 |
| **Volume (ml)** | 50 | 50 | 50 | 50 | 50 |
| **Number of tubes** | 10 | 3 | 2 | 2 | 1 |
| **Food per larvae (mg/larvae/feed)** | 0.55 | 0.55 | 0.55 | 0.55 | 0.55 |
| **Food per tube (mg/tube/feed)** | 3.3 | 8.8 | 15.4 | 19.8 | 27.5 |
| **Ratios** | 1:0, 0:1, 1:1, 2:1, 1:2, 3:4, 4:3, 3:1, 1:3 |  |  |  |  |

A pilot study (see Appendix A) demonstrated that 0.55 mg of liver powder delivered on day 0, 2 and 4 post-hatching supported the successful development of an individual larvae of either species in isolation. Liver powder was delivered as a dilution, mixed so that 1ml of food solution contained the correct concentration of food per-capita for each density treatment. When a larvae pupated, it was decanted to a separate container with a small amount of water until emergence. We recorded the number of survivors and the development time of these survivors. The species and gender of the emerged adult was then determined. We took the development time of a larvae to be from hatching to pupation. A summary of the experimental procedure is given in the Figure 1.

### Analysis

We analysed the survival and development time data using Bayesian generalised additive (GAM) and generalised linear models (GLM). A total of 2435 larvae were analysed in the survival analysis, and 1489 larvae for development time (as only surviving larvae could be analysed). These statistical models were written and fit in JAGS (v4.3) (Plummer, 2003), a Bayesian inference package for the statistical software R (v3.4.2) (R Core Team, 2012). All models mentioned hereafter were run across four chains for a minimum of 2 × 10^5^ iterations, with every second sample discarded to reduce autocorrelation. Chains were continued for additional iterations if they had not yet converged (until the potential scale reduction factor was < 1.02). For all models, parameters were given appropriately diffuse priors centred on zero, unless the pilot studies could be in some way informative (see appendix table 4).

#### Survival

The survival data were modelled as either a binomial distribution or hierarchical beta-binomial distribution. The latter was included after initial inspection of the data showed that the proportions of larvae surviving in tubes undergoing the same density treatment varied. A beta distribution accounts for this variance, and could better explain the data. A binomial regression would model the number of survivors (*S*) from the total number (*N*) in the *i^th^* tube as the outcome of a binomial distribution with a probability *p*

**Figure. 1:**
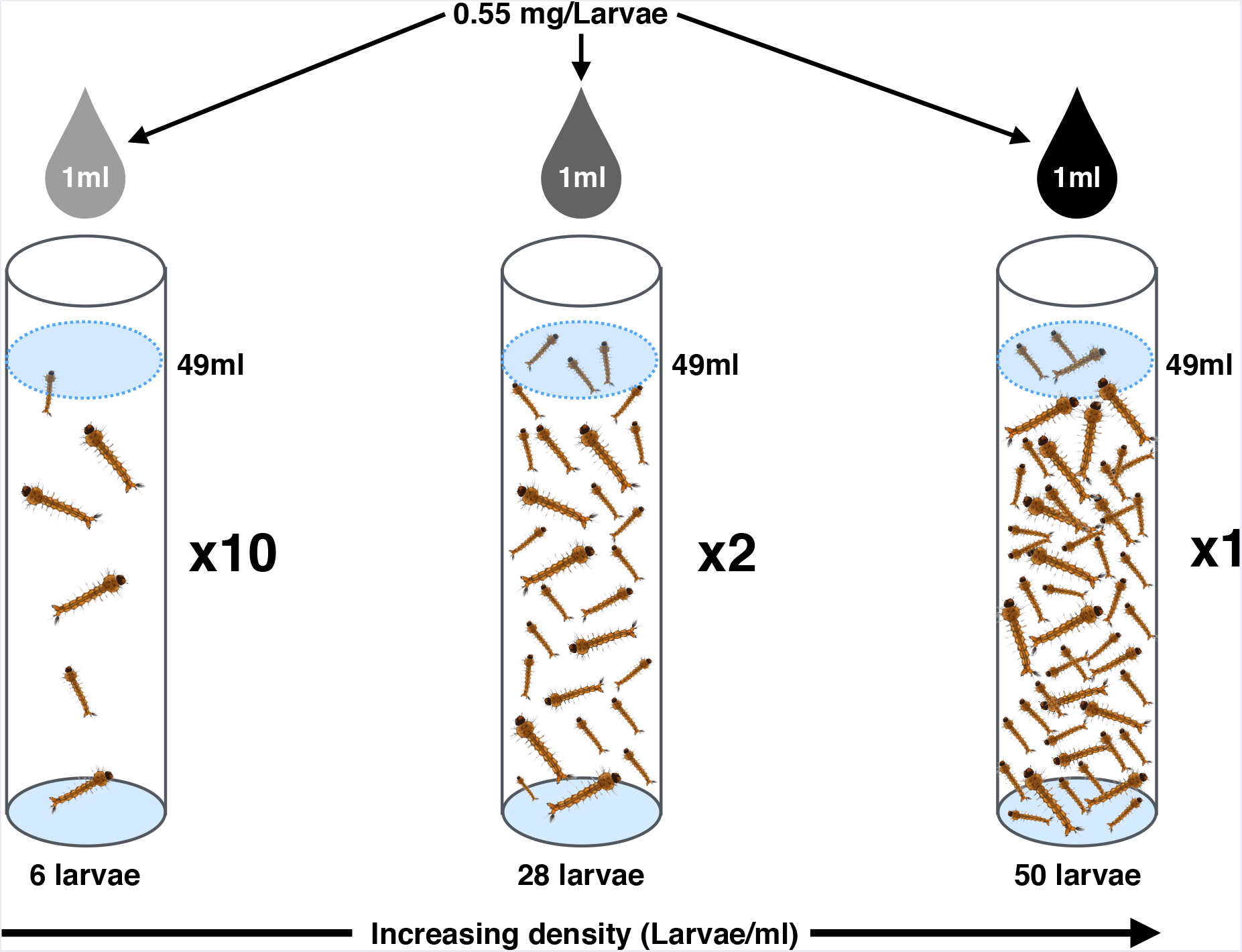
Summary schematic of the experimental design and procedure, shown for a 1:0 species ratio and 3 of the 5 density treatments tested. Falcon tubes were filled with reverse-osmosis water to 49ml, then made up with a final 1ml of a food dilution. Food dilutions were mixed so that 1ml held 0.55mg per larvae in suspension for each treatment (therefore a higher concentration was needed to cater for more larvae in higher density treatments). A further 1ml was added on day 2 and 4 of development. A number of larvae corresponding to one of the densities (6, 16, 28, 38 and 50 larvae for 0.12, 0.32, 0.56, 0.76 and 1 larvae per ml) were hatched and added to the falcons using a pipette. As the low density treatment contained only 6 individuals and the high density 50, it was necessary to repeat these tubes 10 times to achieve a similar sample size. All density treatments were repeated so that there were around 50 individuals experiencing each treatment. Pupated individuals were counted every day, and were removed and decanted to a small water filled tube until emergence. The emerged adults were sexed and identified to species level (for mixed species experiments). This procedure was repeated twice for each of the species ratios in Table 1.

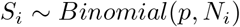

The probability p is written as a function of explanatory variables (say a vector of predictors *X*) (*p* = *f* (*X*)). A logit-link was used to bound p between zero and one (*logit*(*M*) = *f* (*X*)).

In the case of the beta-binomial distribution, the probability is given by a beta distribution with with two shape parameters, *s*_1_ and *s*_2_

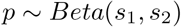

This beta distribution is re-parametrised in terms of the mode of the distribution, for the following two reasons. First, neither *s*_1_ or *s*_2_ describe a useful property of the beta distribution (e.g. mean, standard deviation) to write as a function of covariates. Second, the mode is a far better description of skewed distributions than other statistics such as the mean, which can be heavily influenced by “long tails”. The re-parametrisation is as follows, with the mode *M* and a certainty parameter *θ* (Kruschke, 2015)

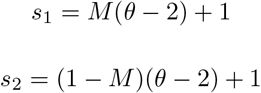

In this case the mode *M* is written as as a function of the covariates (*M* = *f* (*X*)). A logit-link was used to bound the mode between zero and one (*logit*(*M*) = *f* (*X*)).

Exploration of the survival data showed some evidence that the survival probability could be changing as a non-linear function of *Ae. aegypti* and *Ae. albopictus* density, and with some irregularity. Generalised additive models (GAMs) use data-derived splines to fit smoothed non-linear responses to continuous explanatory variables (Wood, 2006). As they are derived from the data, smoothed splines are not constrained in the same way higher order self-interaction terms are in standard linear models. We therefore included smoothed responses in both the binomial and beta-binomial models in addition to the linear terms. GAM terms can be estimated in a Bayesian framework by modelling the smoothing parameters as random effects, where each smoothing parameter comes from a the same normal distribution with an estimated standard deviation (Crainiceanu, Ruppert, and Wand, 2005). We used 5 knots for our splines, one per point on the density scale.

#### Development time

We modelled the development times as a gamma distribution (positive continuous values). As with the beta distribution, the shape and rate parameters of the gamma distribution do not describe any useful property to model as a function of the covariates. Additionally, when close to zero, the gamma distribution can be skewed. We therefore re-expressed the shape (*k*) and rate (*r*) parameters of the gamma distribution in terms of the mode, *M*, and standard deviation, σ (Kruschke, 2015)

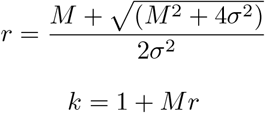

where the mode is written as a function of explanatory variables with a log-link, to ensure the mode is positive (*log*(*M*) = *f* (*X*)). The form of *f* (*X*) was assumed to be a standard linear model, as there was no evidence of non-linearity beyond that afforded by the link function.

### Selecting predictors and model structures

#### Gibbs variable selection

Thoroughly exploring all biologically relevant combinations of predictors can be extremely time consuming, particularly when dealing with multiple categorical interactions. In our case, development time can be explained by two categorical variables, species and sex, and two continuous variables, *Ae. aegypti* and *Ae. albopictus* density. To test the full model space, it would be necessary to explore all second and third order interactions between these terms (this is similarly the case for survival, less the sex variable). We therefore opted to use Gibbs variable selection (GVS), which combines variable selection and model estimation into the same step (Tenan et al., 2014).

In brief, GVS adds a set of binary variables to the model, which are capable of turning individual parameters on and off during optimisation. As an example, we could write a generic response variable *y* as being predicted by p predictors. This model could be written in matrix form, using a vector of parameters β (length p) and a design matrix, **X**

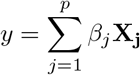

In the case of GVS, a column vector of binary indicator variables, *γ* is added, where *ϒ* ∊ {0,1}. Each each element of *γ* corresponds to a parameter in *β*.

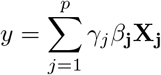

The state of each of element of *ϒ* is denotes whether predictor should to be included in the model (1), or excluded (0). Each element of *γ* is assigned an unbiased 50:50 *Bernoulli* prior (i.e. there is a prior probability of 0.5 of any one parameter being included). The frequency with which combinations of model variables are activated can then interpreted as the preference for including predictors.

In our case, this procedure is complicated by the inclusion of two and three way categorical interactions with hereditary constraints (the constituent, lower-order terms of an interaction must also be included in a model). The priors for the *γ* terms of the interactions were therefore written as the multiple of the states of the lower order terms (*γ_i_* ~ *Bernoulli*(*γ_j_π_i_*), where *j* denoted a lower order term on which the higher order term, *i*, depends). This ensured that the state of the interaction indicator could only be one (*γ_i_* = 1) if the lower order terms were also active (*γ_j_* = 1). This does however make higher order interactions inherently less likely to be included (Kruschke, 2015), though this is not necessarily undesirable for high order interactions terms should only be included with strong support.

Our implementation of GVS also makes use of pseudo priors, as suggested by Dellaportas, Forster, and Ntzoufras (2000). Pseudo priors are priors which are do not affect the posterior distribution of the estimated parameters, instead they are designed to improve the performance of the MCMC sampler itself. Pseudo priors were taken from an initial model run where all indicator variables were set to 1. This model was therefore the “full model”, with a posterior estimates obtained for each parameter. These posteriors are then used as pseudo priors, where they are only active when *γ_j_* = 0. Otherwise (i.e. when the parameter is active and being optimised) the standard prior is used. This can be written, for a normally distributed parameter, with the mean and standard deviation either given as pseudo priors from an initial model run (*μ*̅ and *σ*̅) or as the actual priors (*μ* and *σ*) depending on the state of *γ*

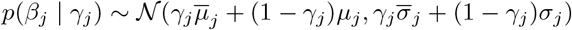

Where *β_j_* is an element of the parameter vector, and *γ_j_* the corresponding element of the vector of binary variables. The motivation for using pseudo priors is to encourage the sampler to frequently turn model parameters on and off, as only if the full model space is thoroughly explored can adequate parameter estimates be obtained, and the inference be trusted. Pseudo priors achieve this by sampling from the posterior estimate of the a parameter (the pseudo prior) when it is inactive, encouraging it to switch the parameter on (the posterior will clearly be more likely than the prior). Turning the parameter back on, the prior returns to the true values and the model is estimated as normal.

#### Product space method

GVS is useful for selecting predictors as the full model space does not need to be explicitly specified, saving time and effort. It is however not suited to the comparison of model different model *structures* and *distributions* (such as the binomial and beta-binomial being compared for the survival data). Bayes factor can be used to compare the likelihood of the data under two statistical models, even when they are not nested. Models are assessed on the prior-weighted average of how well they fit the data, over the whole parameter space (Lodewyckx et al., 2011). Complexity is implicitly penalised by this averaging, as models with many poorly fitting parameters will preform worse in aggregate than a model with a few well fitting parameters. The product space method (PSM) is a way of obtaining a Bayes factor to compare between model structures.

Lodewyckx et al. (2011) explain how the PSM can yield the Bayes factor for multiple models. They suggest that all models should be fit simultaneously as part of an aggregate model, with a stochastic categorical distribution coded to select which model is active. The frequencies with which each model is selected to explain the data is then a measure of the Bayes factor. The use of pseudo priors is suggested in order to promote switching, and they operate in the same way as those in GVS. Pseudo priors are obtained by optimising each model separately, obtaining posteriors for each. As before, the pseudo priors do not affect the likelihood, and there therefore do not influence the posterior directly. They are included solely to improve the performance of the sampler itself, and promote frequent switches. Other checks recommended in (Lodewyckx et al., 2011) were also carried out to ensure that Bayes factor estimates were robust. A preferred model was chosen based on the thresholds given in Raftery (1995).

All parameter estimates and predictions are presented as medians, with 95% highest density intervals (HDI’s) as a measure of uncertainly. We elected to use HDI’s as they are less misleading than symmetric quantiles when describing skewed sample distributions, as can be produced by MCMC methods (Kruschke, 2015). HDI’s correspond to a range across which 95% of the probability mass of a sample falls, and is therefore an intuitive metric of uncertainty.

## Results

### Survival

GVS selection yielded optimal predictor combinations for the two candidate model structures (the binomial and the beta-binomial). These GVS results are reported in full in appendix Tables 5 and 6. The two best models from the GVS were then compared using the product space method. This yielded a log Bayes factor strongly in favour of the beta-binomial model (*log*(*BF*_12_) = −36.54). The hierarchical beta-binomial model was therefore selected as the best model.

This model included a species-specific intercept, a species-specific response to *Ae. aegypti* density and a species-agnostic response to *Ae. albopictus* density. Notably, none of the non-linear responses were included in the final model, implying that the process could adequately be captured by a linear response. This combination of predictors was selected in 33.51% of 4 × 10^6^ iterations, with the next nearest parameter combination at 24.20% (see Table 6 for full GVS results). Parameters estimates for this model are given in Table 2 and shown in Appendix C Figure 8.

In the lowest density single-species treatments, *Ae. aegypti* was 6.58% [0.76, 15.89] more likely to survive than *Ae. albopictus. Ae. albopictus* survivorship declined in response to *Ae. aegypti* density, and did so more severely than *Ae. aegypti*. Across the density range, the estimated intraspecifc response of *Ae. aegypti* was a reduction in survival probability of 17.36% [5.69, 29.33], while the interspecifc response of *Ae. albopictus* was a reduction of 37.99% [21.67, 52.01]. The species did not differ in how they responded to *Ae. albopictus* density, with the intraspecific response of *Ae. albopictus* and interspecific response of *Ae. aegypti* being a 22.81% [10.98, 34.65] reduction in survival probability. Figure 2 shows the effect of conspecific density on the survival probability of both species (intraspecific competition), while Figure 3 shows the effect of both conspecific and heterospecific density (intraspecific *and* intraspecific).

### Development Time

The most frequently selected model during GVS was one including a species- and sex-specific adjustment to the intercept as well as species-specific response to *Ae. aegypti* density, *Ae. albopictus* density and the *interaction* between these two densities. This model was selected in 22.92% of 4 × 10^6^ iterations, with the next nearest model at 21.95% (8.70% after that). The full GVS results are given in appendix Table 7. This next nearest model excluded the interaction term and the species-specific response to either density, meaning it was nested in the former. As these two models were selected with a comparable frequency and therefore had similar support, we compared these two models using the product space method to further assess the performance of each model. The Bayes factor supported the more complex model (*log*(*BF*_12_) = −0.795), which included the species specific effects and interaction.

Parameter estimates are given in Table 3 (and shown in appendix Figure 9). The development times of *Ae. aegypti* in the lowest density treatment were 15.52% [11.84, 19.30] faster than those of *Ae. albopictus*. Females of both species took 7.85% [5.86, 10.01] longer to develop than males. Increasing densities of conspecifics led to faster development times for both species, but far less so for *Ae. aegypti*. Across the density range intraspecific competition reduced development times by 8.37 % [4.00, 12.61] for *Ae. aegypti* and 16.14 % [11.19, 20.72] for *Ae. albopictus*. Interspecific competition reduced development times by 10.86% [0.53, 20.30] for *Ae. aegypti* and 25.39% [14.98, 34.92] for *Ae. albopictus*. The intraspecific effects are illustrated in Figure 4, and the combined intra- and interspecific effects in Figure 5.

**Figure. 2:**
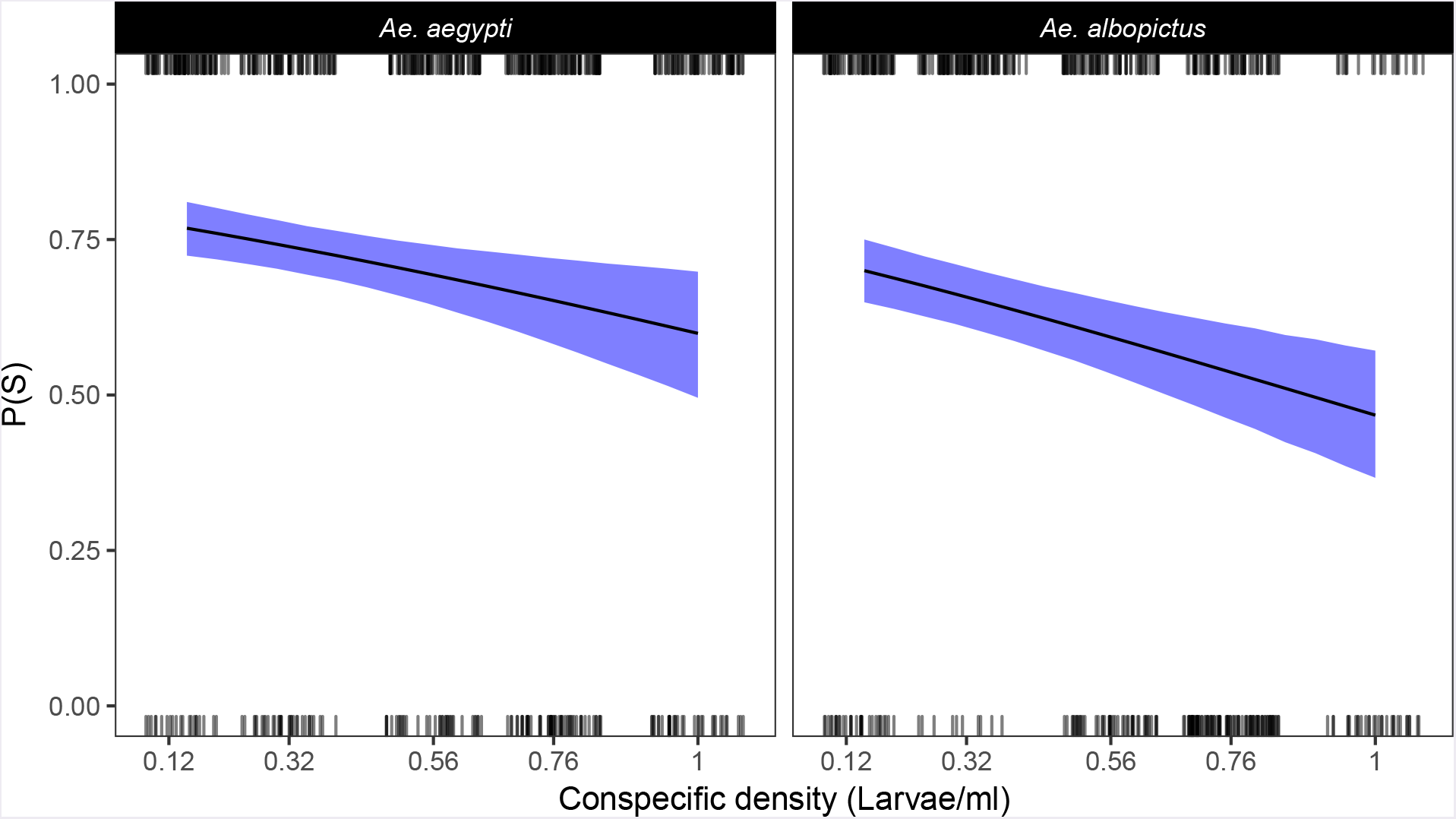
Model fit for the beta-binomial model of the survival data. Data are shown as jittered tick marks on the floor (deaths) and ceiling (survivors). Both species respond linearly to conspecific density (as no nonlinear generalised additive model terms were retained through GVS). *Ae. albopictus* responds more strongly to conspecific density than *Ae. aegypti* (17.36% [5.69, 29.33] reduction versus 22.81% [10.98, 34.65]).

**Figure. 3:**
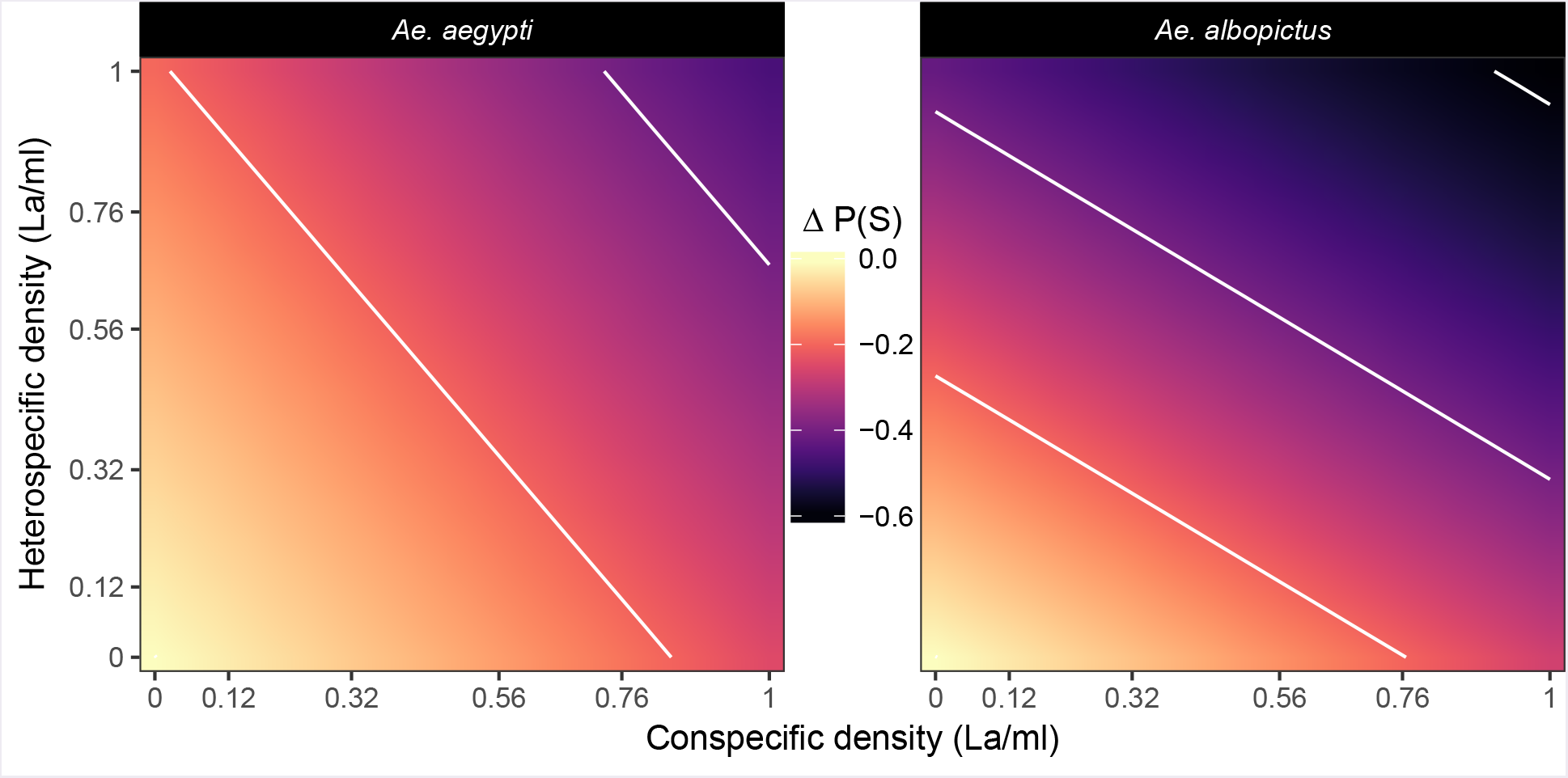
Changes in the survivorship predicted by the beta-binomial survival model across the combinations of density conditions outlined in Table 1. The intraspecific responses run along the x-axis and the interspecifc response along the y-axis. Off-axis values are the additive combinations of inter- and intraspecific competition. Values are expressed as the difference from the baseline survival probability (Δ P(S)), which is the probability associated with the bottom left hand corner in each panel. Contour lines delineate 20% decreases in survival probability, with colder colours indicating decreases in survival.

**Table 2:** Parameter estimates for the beta-binomial model of the survival data, after GVS. Values are quoted as the medians, with the 95% highest density intervals also given. Values are given for each species if there was species specific response included in the final model. Note these values are on a logit scale. Note that the value for *Ae. albopictus* density is the intraspecific density for *Ae. albopictus* and interspecific competition for *Ae. aegypti*.

| Parameter | Median | Lower 95% - Upper 95% |
| --- | --- | --- |
| ***Ae. aegypti*** |  |  |
| Intercept | 1.34 | 1.04 to 1.65 |
| Intraspecific | -0.94 | -1.55 to -0.33 |
| ***Ae. albopictus*** |  |  |
| Intercept | 1.02 | 0.73 to 1.31 |
| Interspecific | -1.83 | -2.71 to -0.96 |
| **Both species** |  |  |
| *Ae. albopictus* density | -1.15 | -1.70 to -0.60 |
| $\theta$ (certainty parameter) | 22.94 | 16.57 to 30.86 |

Interestingly, the interactive effect of density for *Ae. aegypti* was estimated as being tightly centred on zero (−0.01 [-0.06, 0.03], log scale), meaning that it did not respond differently to combined higher densities of both species. In contrast, the interactive response of *Ae. albopictus* was strongly positive (0.11 [0.06, 0.16], log scale). The model predicted that at the maximum *combined* densities of both species, surviving *Ae. albopictus* would take 59.11 % [11.03, 109.54] *longer* to develop compared to the lowest density treatment. This is contrary to the faster development time observed when responding to conspecific and heterospecific density in isolation. The interactive effect is observable in Figure 5, towards the back right corner of the *Ae. albopictus* panels.

**Figure. 4:**
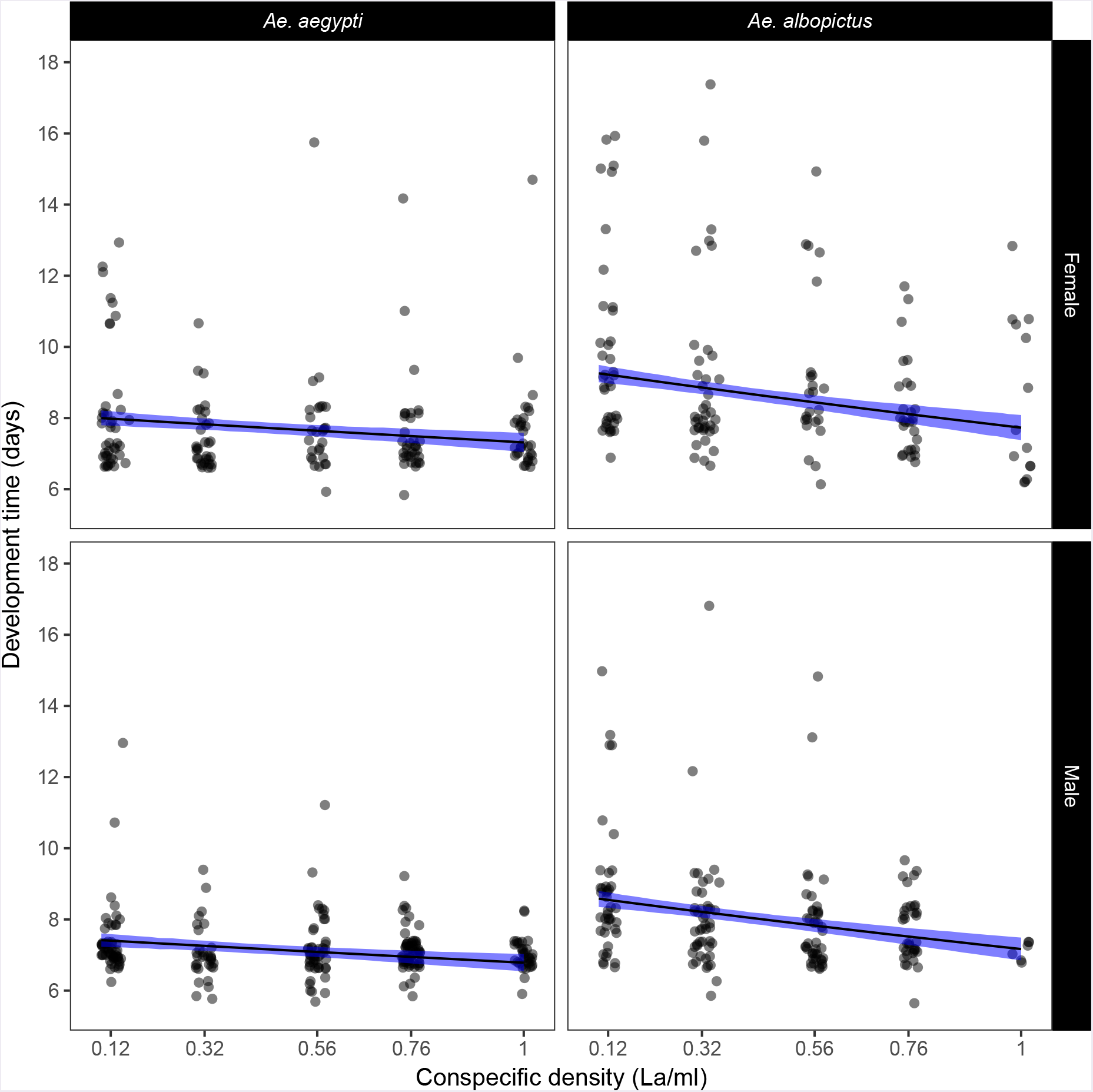
Model fit for the gamma-distributed model of the development time data. This figure shows intraspecific competition for both species. Evident is the slower baseline development time of *Ae. albopictus* (15.52 % [11.84, 19.30] slower), and marginally slower development of females for both species (7.86 % [5.86, 10.01]). Note that there is no sex-specific gradient, only an adjustment to the intercept. Across the range of densities, development times are reduced by 8.37 % [4.00, 12.61] for *Ae. aegypti* and by 16.14 % [11.19, 20.72] for *Ae. albopictus*.

**Figure. 5:**
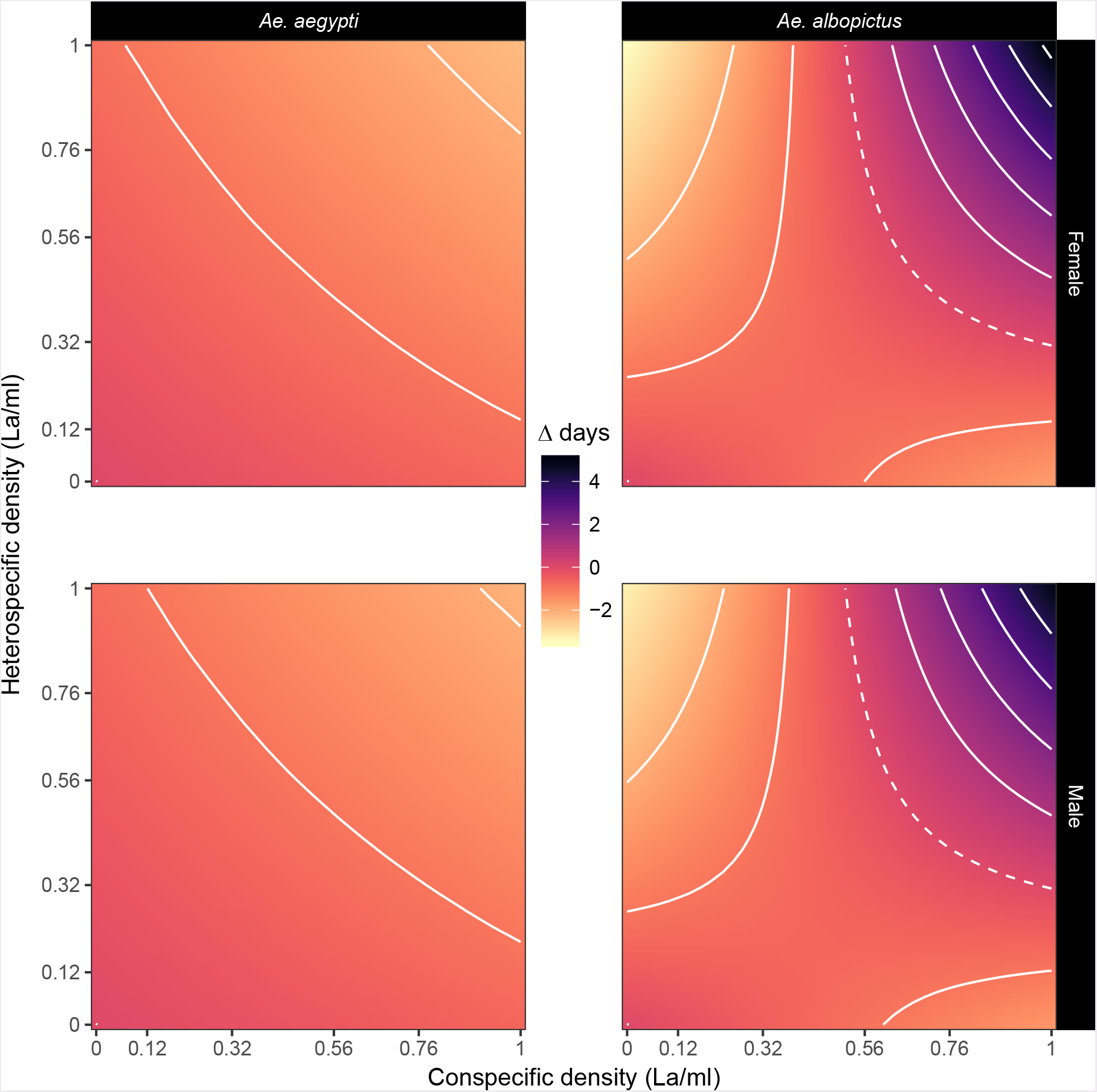
Changes in the predicted development time (Δ days) across the combinations of density conditions outlined in table 1. Within-species competition is shown along the x-axis and between-species on the y-axis. Values are expressed as the differences from the intercept development time in each panel (the development time associated with the bottom left hand corner of each panel). Colours warmer than the zero mark in the legend indicate decreases in development time, while colder colours indicate increases in development time (compared to the bottom left-hand value). Contour lines delineate changes of 2 days, with the dashed contour the zero mark. There is notably little change in the development time of *Ae. aegypti* across the surface (smaller effect sizes), with no interactive effects of density. In contrast, *Ae. albopictus* speeds up it’s development time in response to hetero- and conspecific density, albeit to a greater extent for heterospecifics. However, the positive interactive effects of density on *Ae. albopictus* development time means the model predicts slower development when densities of conspecifics *and* heterospecifics are high. Differences between males and females are minor, and determined only by the shift in the intercept (there were no sex-specific responses to density).

**Table 3:** Parameter estimates for the gamma-distributed GLM for development time, following GVS and co- marison by the PSM. Values are quoted as the medians, with the 95% highest density intervals also given. Values are given for each species where there are species specific responses. All values are on a log scale. The predictor for the interaction term was multiplied by 10 to keep it on the same scale as the two independent densities, so the effect size is directly comparable. f symbols denote parameters with intervals that cross zero. These have been retained due to hereditary constraints in predictor selection.

| Parameter | Median | Lower 95% - Upper 95% |
| --- | --- | --- |
| ***Ae. aegypti* males** |  |  |
| Intercept | 2.01 | 1.98 to 2.04 |
| Intraspecific density | -0.10 | -0.15 to -0.05 |
| Interspecific density † | -0.12 | -0.29 to 0.05 |
| Density interaction † | -0.01 | -0.06 to 0.03 |
| ***Ae. albopictus* males** |  |  |
| Intercept | 2.17 | 2.14 to 2.20 |
| Intraspecific density | -0.20 | -0.27 to -0.14 |
| Interspecific density | -0.47 | -0.68 to -0.26 |
| Density interaction | 0.11 | 0.06 to 0.16 |
| **Both species** |  |  |
| Sex (female) | 0.08 | 0.06 to 0.10 |
| $\sigma$ (standard deviation) | 1.48 | 1.42 to 1.53 |

## Discussion

As vectors of diseases such as dengue and Zika, the ecology of *Aedes* mosquitoes is of the utmost importance for predicting vector occurrence, disease incidence and the efficacy of interventions. Our study aimed to quantify the effects of inter- and intraspecifc competition on the survivorship and development time of larvae under a particular per-capita feeding regime. We find that *Ae. aegypti* suffered intra- and interspecific competition to a lesser extent than *Ae. albopictus*. Indeed, effect sizes were very small for *Ae. aegypti* (particularly for development time) whereas *Ae. albopictus* suffered higher mortality and took longer to develop when in high conspecific and heterospecific densities. However, the fact we observe any effect is of note, as the larvae were fed per-capita, and at a level which *could* have supported the successful development of all larvae.

The higher baseline survivorship of *Ae. aegypti* is consistent with historic lab studies (Macdonald, 1962) and studies using a similar food resource (Barrera, 1996), but not other studies using other food types (Juliano, Lounibos, and O’Meara, 2004). There was no support for a species-specific response of survivorship to *Ae. albopictus* density, indicating that the strength of intraspecific competition in *Ae. albopictus* is equal to the effect of interspecific competition on *Ae. aegypti*. The same is not true for the response to *Ae. aegypti* density, where there is a greater effect of interspecific competition on *Ae. albopictus* than intraspecific competition on *Ae. aegypti*. The findings for survivorship run somewhat contrary to other competition studies, where the effects of on interspecific competition are usually greater for *Ae. aegypti* (Juliano, Lounibos, and O’Meara, 2004; Reiskin and Lounibos, 2009) or neutral (Black et al., 1989). This could be explained by the use of liver powder as our choice of larval resource, as this has been known to favour *Ae. aegypti* (Barrera, 1996). Alternatively, this could be the result of strain-specific properties (Dye, 1984a).

The superiority of the beta-binomial model structure in explaining the survivorship data tells us that an additional second-level process was required to account for the variance in larval survival between tube replicates. This variance could either be “true” demographic variance in survival or “error” variance caused by our experimental setup across tubes and replicates (miscounts of larvae, food dilutions). Should it be the former, this is indicative of interesting features of larval demographic variation. In low-density treatments there were fewer individuals, a difference which we accounted for by repeating these tubes (so that the numbers per treatment were the same). However, we could not control for the demographic make-up of mosquitoes in each tubes, and therefore the dynamics in low-density tubes will be effected by the sample of larval phenotypes expressed in this relatively small sample of larvae. For instance, if each larvae has a certain capacity to respond to conspecifc and heterospecifc density (e.g. increased resource uptake), then in a small cohort a *single* larva capable of outperforming the others would have a profound effect on the density effects observed in the tube. Individual phenotypes are known to generate the population dynamics we observe (Sumpter and Broomhead, 2001), and it is of note that in smaller cohorts of mosquitoes variation in processes such as survival could shape the observed patterns of persistence and abundance. This evidence points to the importance of demographic processes in the overall dynamics of mosquito populations.

Across the combinations of densities, surviving *Ae. aegypti* larvae developed more quickly than *Ae. albopictus*. In response to conspecific density, the surviving larvae of both species were those that developed quickly. This suggests that the surviving larvae either accrued the necessary resources to pupate faster than competitors, or had a lower resource threshold for pupation. *Ae. aegypti* development times changed only marginally, whereas *Ae. albopictus* did so to a far greater extent.

Of most interest is the observation that *Ae. albopictus* larvae which survived in treatments with high densities of conspecifics *and* heterospecifics developed more slowly. This is the opposite of its response to intraspecific competition, where it sped up development time. It could be the case that slowly developing individuals are able to cannibalise deceased larvae which break down over time (dead larvae were not removed during counts). It is known that invertebrate carcasses can be a food source for larva (Daugherty, Alto, and Juliano, 2000), so perhaps some *Ae. albopictus* larvae are “playing the long-game”, and benefiting from the delayed release of this additional resource. The longest development times for *Ae. albopictus* also occur under conditions where the mortality of both species is highest, meaning the number of carcasses would also be highest. This fits with evidence that *Ae. albopictus* is better able to survive periods of starvation (Barrera, 1996). While the chemical retardant hypothesis (Moore and Fisher, 1969; Moore and Whitacre, 1972) has received little recent attention, it is worth mentioning that this mechanism could also account for the delayed development time observed in *Ae. albopictus*.

Our study is novel in that our feeding regime was adjusted for each density treatment so that the per-capita resource availability was the same. From our pilot studies we knew that this amount was sufficient for a larvae of either species to pupate. Despite the larvae of both species feasibly being able to pupate on the available resources, many did not, especially for *Ae. albopictus* at high densities. The implication is that non-pupating larvae were not getting the threshold resources required to become pupae. This could be attributed to con- and heterospecifics taking more than the “allotted” 0.55mg per larvae. Interestingly, this points to the rate and degree of resource uptake being mediated by the density of other larvae - both of the same and different species. It would seem that *Ae. aegypti* is better able to alter these resource seeking/uptake parameters than *Ae. albopictus*, as it suffers fewer deaths as a result of competition. Had our study provided even more food per-capita, it is possible that the effects of competition would have been ameliorated. This is because there may come a level of food availability where even significant changes in the rate and extent of uptake by larvae would not be enough to inhibit competitors. Repeating this experiment at a greater range of per-capita feeding regimes is certainly an avenue worth pursuing.

This is therefore a manifestation of contest competition, whereby the uptake of resources by the faster developing larvae results in the death of slower developing larvae who never gain the requisite resources for pupation. In contrast, if all larvae expressed the same resource seeking and accruing behaviours that they had in isolation, there would have been sufficient food for all to pupate.

It is noteworthy that in a single-species example (using only *Ae. aegypti*), Heath et al. (*In review*) did not find an effect of conspecific density on either survival and development time using this exact feeding regime. The only difference between the experiments was the type and volume of containers used to rear the larvae. This may allude to an effect of habitat volume and surface area, with individuals potentially competing for space, and access to the surface to breathe. Smaller containers with smaller surface areas may amplify the behavioural responses, leading to the outcomes described above. Such mechanisms have been suggested in the past (Dye, 1984b), and further emphasise the extent to which habitat properties can alter competitive outcomes.

## Conclusion

When seeking a consensus from competition experiments between *Ae. aegypti* and *Ae. albopictus*, Juliano (2009) and others found that competition was context dependent. Outcomes varied across gradients of habitat resource availability, temperature and constancy. It was therefore never the case that any one study could *fully* explain the competition between these two species. However, our study contributes a unique insight into the consequences of competition under this specific feeding regime. The results show how these larvae could potentially alter the rate and extent of resource uptake to the detriment of their peers. We also provide evidence of an interactive effect on *Ae. albopictus* development time, which we suggest could be mediated by mechanisms other than resource uptake, such as cannibalism or chemical interference. Our model selection revealed that our survival data were overdispersed, which highlights the importance of demographic processes in driving variation in population-level processes. We add to the broader understanding of how within and between species competition can be driven by both resource mediated and resource independent processes, and give insight into the degree to which even relatively simple organisms can display a diverse range of responses to increased densities of competitors.

## Acknowledgements

We would like to thank Jo Stoner, Sanjay Basu and Derek Au, who supported RSP and KH with the experiments. Comments and advice from Chris Terry were appreciated.

RP was supported by a NERC studentship (NE/L002612/1) and is a CASE student with the Pirbright Institute. KH was supported by a BBSRC studentship (BB/M011224/1).

## Author contributions

RSP and KH designed the experiments and developed the concept and ideas, with advice from MBB and AJW. RSP and KH carried out the experiments. RSP designed and conduced the analysis, and wrote the manuscript. KH, MBB and AJW reviewed the manuscript, and contributed to the final document.

## A Pilot study

In order to choose an appropriate feeding regime for the main experiments, it was necessary to find the baseline per-capita nutritional requirements of both species. We therefore explored, on a per-capita basis, how much liver powder was required for a larvae to successfully develop.

### Procedure

Eggs of each species were vacuum hatched for approximately 1.5 hours in tap water, which ensured all larvae were the same age. The wells of a 6-well-plate were filled with 9ml RO filtered water. A single larvae was placed into each well of a six well plate. A dilution of liver powder corresponding to 0.05, 0.2, 0.35, 0.5, 0.65 and 0.8 mg per larvae was added to each well, bringing the total volume to 10ml. The larvae were fed again with 1ml of this solution on the 2nd and 4th day after hatching (evaporation was thought to maintain the volume at ~ 10 ml). Each day the larvae were checked to see if they had pupated or died. Larvae were given a three weeks to develop, after which time they were assumed dead. This process was repeated for each species, across 3 replicates.

A binomially distributed logit-link generalised linear model was used to analyse the survival data, and a step-wise down model selection procedure using log-likelihood ratio tests (LRT) was used to determine the optimal set of predictors.

### Results

Food treatment had a significant (LRT: Δ*df* = 1, *p* = 2.1 × 10^−16^) positive effect (0.0143 ± 0.0025, logit scale) on survivorship, but the effect did not significantly differ between species (LRT: Δ*df* = 1, *p* = 0.1299). The relationship is shown in Figure 6.

### Conclusions

Both species responded to the availability to the liver power in the same way. The estimated probability of survival asymptotically approached 1 from approximately 0.5mg/La. We therefore elected to set our feeding regime at 0.55mg/La, to ensure that there were adequate food resources to maintain all larvae in the population.

**Figure. 6:**
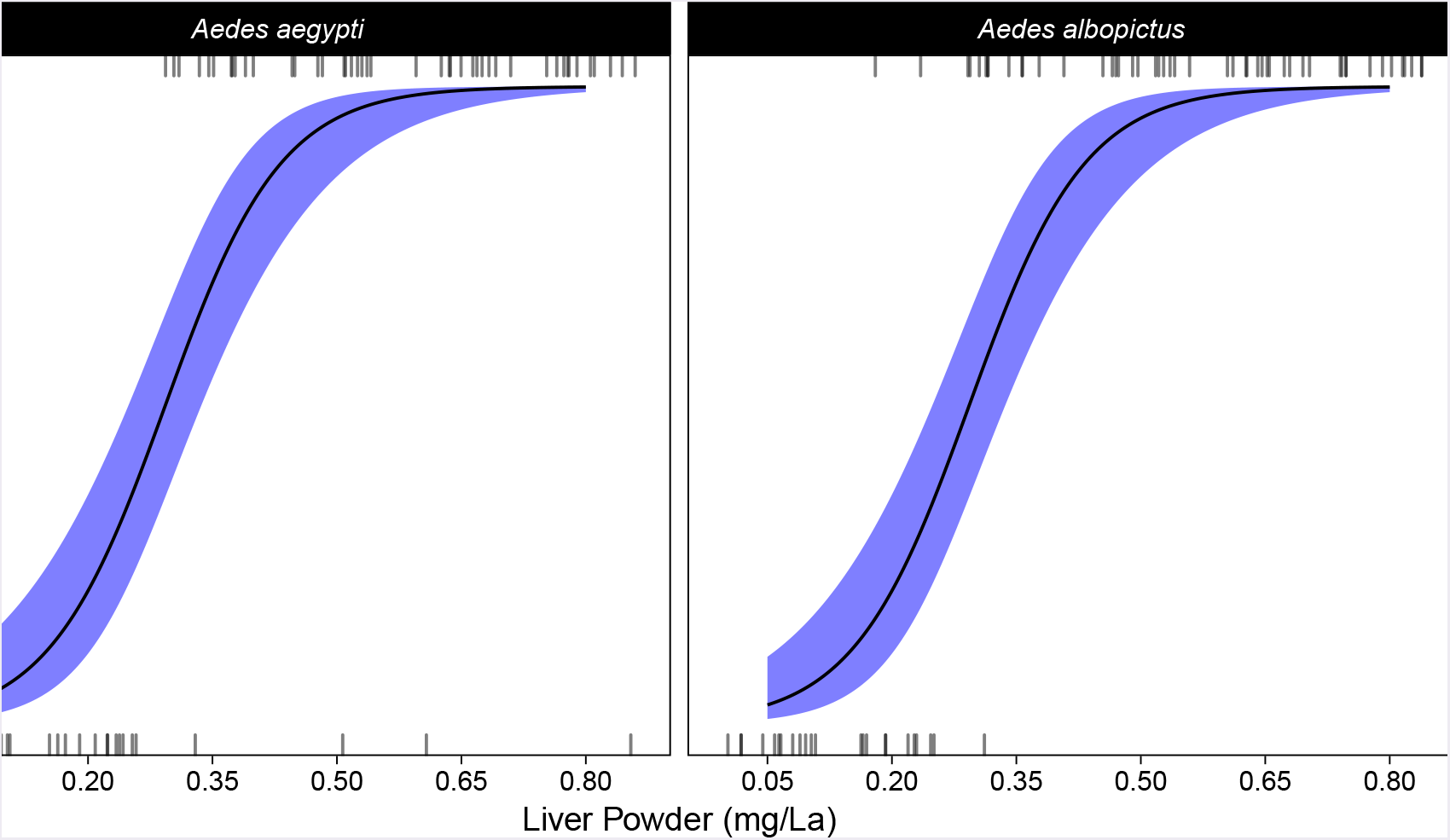
Experimental data for the pilot studies examining the development time and survivorship of *Ae. aegypti* and *Ae. albopictus*. Points show the per-replicate proportion of larvae that survived each food treatment, while the rugs show the raw survival data. Note that there is a positive effect of food availability on survivorship, that it is consistent across both species.

## B Priors

**Table 4:** Priors used for fitting the binomial, beta binomial and development time models. Both intercepts were taken from pilot runs of the experiment, as was the standard deviation for development time.

| Survival Models |  | Development Time Model |  |
| --- | --- | --- | --- |
| Variable(s) | Distribution | Variable(s) | Distribution |
| Intercept | $\mathcal{N}(2.7, 1)$ | Intercept | $\mathcal{N}(2.079, 1)$ |
| Linear predictors | $\mathcal{N}(0, 1)$ | Linear predictors | $\mathcal{N}(0, 0.3)$ |
| Smoothing parameters | $\Gamma(1, 0.001)$ | Standard deviation | $\Gamma(1.28, 0.23)$ |
| Certainty, $\theta$ (beta-binomial) | $\Gamma(1, 1000)$ | - | - |

## C Gibbs Variable Selection (GVS)

Results for the GVS procedure are reported for the two survivorship model structures (binomial in table 5 and beta-binomial in table 6), and the gamma model of development times (table 7). The frequencies with which predictor combinations are reported, along with the absolute number of iterations. Parameter combinations selected less than 1% of the iterations are not included in the tables. The fits corresponding to each of the best fitting model are given in Figures 7 and 8 for the two survival models, and 9 for the development time model.

**Table 5:** Frequencies with which combinations of predictors were selected for the binomial model during GVS. *N* denotes for how many samples particular variable combinations were active, and the percentages calculated as *N* divided by the total number of iterations (4 × 10^6^). Int is the intercept, *Sp* the species, *D_aeg_* the density of *Ae. aegypti* and D_alb_ the density of *Ae. albopictus*. Interactions between variables are denoted by :, while *f* () denotes that the term is a smooth function of the predictor in brackets. Each smooth functions had 5 knots. The most frequently selected model is given in row 1, and included a species-specific intercept, a linear species specific response to *Ae. albopictus* density and a linear and smooth term for *Ae. aegypti* density.

| Rank | Predictor combinations | $N$ | % |
| --- | --- | --- | --- |
| 1 | $Int + Sp + D_{aeg} + D_{alb} + Sp : D_{alb} + f(D_{aeg})$ | 633723 | 15.84 |
| 2 | $Int + Sp + D_{aeg} + Sp : D_{aeg} + D_{alb} + Sp : D_{alb} + f(D_{aeg})$ | 402459 | 10.06 |
| 3 | $Int + Sp + D_{aeg} + Sp : D_{aeg} + D_{alb} + D_{aeg} : D_{aeg} + f(D_{aeg})$ | 326178 | 8.15 |
| 4 | $Int + Sp + D_{aeg} + Sp : D_{aeg} + D_{alb}$ | 295250 | 7.38 |
| 5 | $Int + Sp + D_{aeg} + Sp : D_{aeg} + D_{alb} + Sp : D_{alb}$ | 240442 | 6.01 |
| 6 | $Int + Sp + D_{aeg} + D_{alb} + D_{aeg} : D_{aeg} + f(D_{aeg})$ | 213651 | 5.34 |
| 7 | $Int + Sp + D_{aeg} + D_{alb} + f(D_{aeg})$ | 212110 | 5.30 |
| 8 | $Int + Sp + D_{aeg} + Sp : D_{aeg} + D_{alb} + Sp : D_{alb} + D_{aeg} : D_{aeg} + f(D_{aeg})$ | 202547 | 5.06 |
| 9 | $Int + Sp + D_{aeg} + D_{alb} + Sp : D_{alb} + D_{aeg} : D_{aeg} + f(D_{aeg})$ | 179147 | 4.48 |
| 10 | $Int + Sp + D_{aeg} + Sp : D_{aeg} + D_{alb} + Sp : D_{alb} + Sp : D_{aeg} : D_{alb} + f(D_{aeg})$ | 173728 | 4.34 |
| 11 | $Int + Sp + D_{aeg} + Sp : D_{aeg} + D_{alb} + f(D_{aeg})$ | 152175 | 3.80 |
| 12 | $Int + Sp + D_{aeg} + D_{alb} + Sp : D_{alb} + Sp : D_{aeg} : D_{alb} + f(D_{aeg})$ | 107058 | 2.68 |
| 13 | $Int + Sp + D_{aeg} + Sp : D_{aeg} + D_{alb} + D_{aeg} : D_{aeg} + Sp : D_{aeg} : D_{alb} + f(D_{aeg})$ | 57074 | 1.43 |
| 14 | $Int + Sp + D_{aeg} + Sp : D_{aeg} + D_{alb} + Sp : D_{alb} + Sp : D_{aeg} : D_{alb}$ | 53330 | 1.33 |
| 15 | $Int + Sp + D_{aeg} + Sp : D_{aeg} + D_{alb} + Sp : D_{aeg} : D_{alb}$ | 53317 | 1.33 |
| 16 | $Int + Sp + D_{aeg} + Sp : D_{aeg} + D_{alb} + Sp : D_{alb} + D_{aeg} : D_{aeg} + Sp : D_{aeg} : D_{alb} + f(D_{aeg})$ | 48099 | 1.20 |
| 17 | $Int + Sp + D_{aeg} + D_{alb} + D_{aeg} : D_{aeg} + Sp : D_{aeg} : D_{alb} + f(D_{aeg})$ | 46015 | 1.15 |

**Figure. 7:**
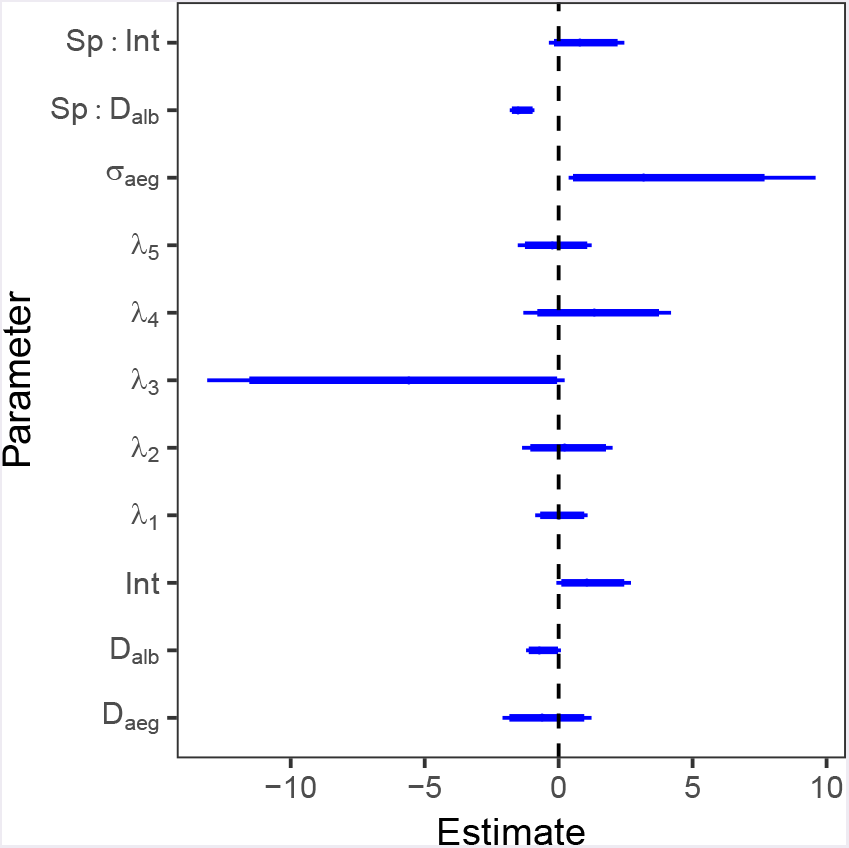
Parameter estimates for the binomial model, after the predictors had been selected by GVS. Dots denote medians, thick lines the 90% HDIs and thin lines the 95% HDIs. In this model, *σ_aúg_* is the standard deviation of the normal distribution used to estimate the smoothing parameters (λ_1-5_) of the 5-knot smooth function of *Ae. aegypti* density (Crainiceanu, Ruppert, and Wand, 2005). Colons denote interactions, “Sp” the intercept change for *Ae. albopictus* and “Int” the intercept. *D_aúg_* and *D_alb_* are the densities of *Ae. aegypti* and *Ae. albopictus* respectively. This model was compared to the beta-binomial model (Figure 8).

**Figure. 8:**
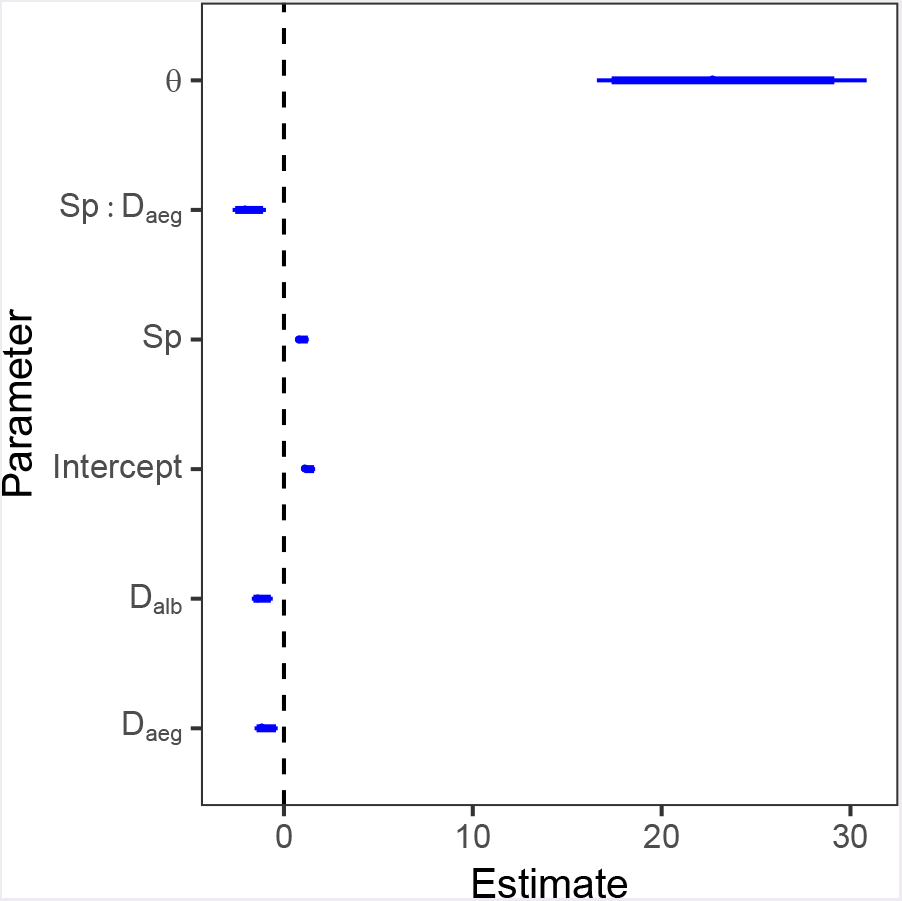
Parameter estimates for the beta-binomial model, after the predictors have been selected by GVS. Dots denote medians, thick lines the 90% HDIs, and thin lines the 95% HDIs. In this model, θ is the certainty parameter, as mentioned in the re-parametrisation of the beta distribution in (Kruschke, 2015). Colons denote interactions, “Sp” the intercept change for *Ae. albopictus* and “Int” the intercept. *D_aúg_* and *D_alb_* are the densities of *Ae. aegypti* and *Ae. albopictus* respectively. This model was compared to the beta-binomial model (Figure 7). These estimates are also given in table 2 in the main text.

Beta-binomial model of larval survival

**Table 6:** Frequencies with which combinations of predictors were selected for the beta-binomial model during GVS. *N* denotes for how many samples particular variable combinations were active, and the percentages calculated as *N* divided by the total number of iterations (4 × 10^6^). Int is the intercept, Sp the species, *D_aúg_* the density of *Ae. aegypti* and *D_alb_* the density of *Ae. albopictus*. Interactions between variables are denoted by :, while *f* () denotes that the term is a smooth function of the predictor in brackets. Each smooth functions had 5 knots. The most frequently selected model is given in row 1, and included a species-specific intercept, a linear species-specific response to *Ae. aegypti* density and a linear *Ae. albopictus* density.

| Rank | Predictor combinations | $N$ | % |
| --- | --- | --- | --- |
| 1 | $Int + Sp + D_{aeg} + Sp : D_{aeg} + D_{alb}$ | 1340494 | 33.51 |
| 2 | $Int + Sp + D_{aeg} + Sp : D_{aeg} + D_{alb} + Sp : D_{alb}$ | 968011 | 24.20 |
| 3 | $Int + Sp + D_{aeg} + D_{alb}$ | 533692 | 13.34 |
| 4 | $Int + Sp + D_{aeg} + D_{alb} + Sp : D_{alb}$ | 355619 | 8.89 |
| 5 | $Int + Sp + D_{aeg} + Sp : D_{aeg} + D_{alb} + D_{aeg} : D_{alb}$ | 184094 | 4.60 |
| 6 | $Int + Sp + D_{aeg} + D_{alb} + Sp : D_{alb} + D_{aeg} : D_{alb}$ | 116732 | 2.92 |
| 7 | $Int + Sp + D_{aeg} + D_{alb} + D_{aeg} : D_{alb}$ | 97547 | 2.44 |
| 8 | $Int + Sp + D_{aeg} + Sp : D_{aeg} + D_{alb} + Sp : D_{alb} + D_{aeg} : D_{alb}$ | 81131 | 2.03 |
| 9 | $Int + Sp + D_{aeg} + D_{alb} + Sp : D_{alb} + f(D_{aeg})$ | 42801 | 1.07 |
| 10 | $Int + D_{aeg} + D_{alb}$ | 41163 | 1.03 |

**Figure. 9:**
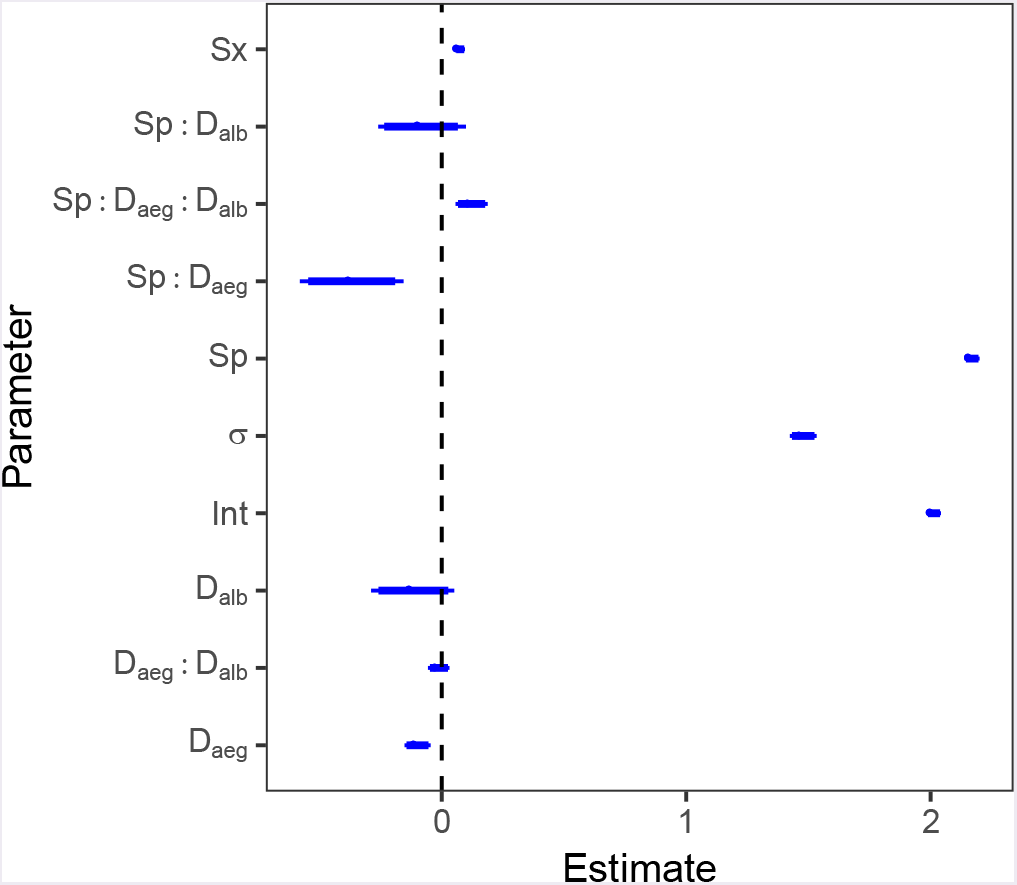
Parameter estimates for the development time model after the predictors have been selected by GVS. Dots denote medians, thick lines the 90% HDIs, and thin lines the 95% HDIs. Colons denote interactions, “Sp” the intercept change for *Ae. albopictus* and “Int” the intercept. *D_aúg_* and D_alb_ are the densities of *Ae. aegypti* and *Ae. albopictus* respectively. These four models were then compared using the PSM to decide on the best structure.

Gamma distributed model of development time

**Table 7:** Frequencies with which combinations of predictors were selected for the gamma GLM during GVS. *N* denotes for how many samples particular variable combinations were active, and the percentages are *N* divided by the total number of iterations (4 × 10^6^). Int is the Intercept, *Sp* the species (categorical), Sx the sex (categorical), *D_aúg_* the density of *Ae. aegypti* and *D_alb_* the density of *Ae. albopictus*. Interactions between variables are denoted by colon. The most selected model is given in the first row, and included a sex-specific intercept, as well as

| Rank | Predictor combinations | $N$ | % |
| --- | --- | --- | --- |
| 1 | $Int + Sp + Sx + D_{aeg} + D_{alb} + D_{aeg} : D_{alb} + Sp : D_{aeg} + Sp : D_{alb} + Sp : D_{aeg} : D_{alb}$ | 916754 | 22.92 |
| 2 | $Int + Sp + Sx + D_{aeg} + D_{alb}$ | 877931 | 21.95 |
| 3 | $Int + Sp + Sx + D_{aeg} + D_{alb} + D_{aeg} : D_{alb} + Sp : D_{aeg} + Sp : D_{alb} + Sx : D_{alb} + Sp : D_{aeg} : D_{alb}$ | 347898 | 8.70 |
| 4 | $Int + Sp + Sx + D_{aeg} + D_{alb} + Sx : D_{alb}$ | 340935 | 8.52 |
| 5 | $Int + Sp + Sx + D_{aeg} + D_{alb} + Sp : D_{aeg}$ | 173395 | 4.33 |
| 6 | $Int + Sp + Sx + D_{aeg} + D_{alb} + D_{aeg} : D_{alb} + Sp : D_{aeg} + Sx : D_{aeg} + Sp : D_{alb} + Sp : D_{aeg} : D_{alb}$ | 154349 | 3.86 |
| 7 | $Int + Sp + Sx + D_{aeg} + D_{alb} + Sp : D_{alb}$ | 153422 | 3.84 |
| 8 | $Int + Sp + Sx + D_{aeg} + D_{alb} + Sx : D_{aeg}$ | 145638 | 3.64 |
| 9 | $Int + Sp + Sx + Sp : Sx + D_{aeg} + D_{alb}$ | 92416 | 2.31 |
| 10 | $Int + Sp + Sx + Sp : Sx + D_{aeg} + D_{alb} + D_{aeg} : D_{alb} + Sp : D_{aeg} + Sp : D_{alb} + Sp : D_{aeg} : D_{alb}$ | 88255 | 2.21 |
| 11 | $Int + Sp + Sx + D_{aeg} + D_{alb} + D_{aeg} : D_{alb}$ | 79092 | 1.98 |
| 12 | $Int + Sp + Sx + D_{aeg} + D_{alb} + Sp : D_{aeg} + Sx : D_{alb}$ | 66902 | 1.67 |
| 13 | $Int + Sp + Sx + D_{aeg} + D_{alb} + Sp : D_{alb} + Sx : D_{alb}$ | 58694 | 1.47 |
| 14 | $Int + Sp + Sx + D_{aeg} + D_{alb} + D_{aeg} : D_{alb} + Sp : D_{alb}$ | 47654 | 1.19 |
| 15 | $Int + Sp + Sx + D_{aeg} + D_{alb} + Sx : D_{aeg} + Sx : D_{alb}$ | 46270 | 1.16 |

**Figure. 10:**
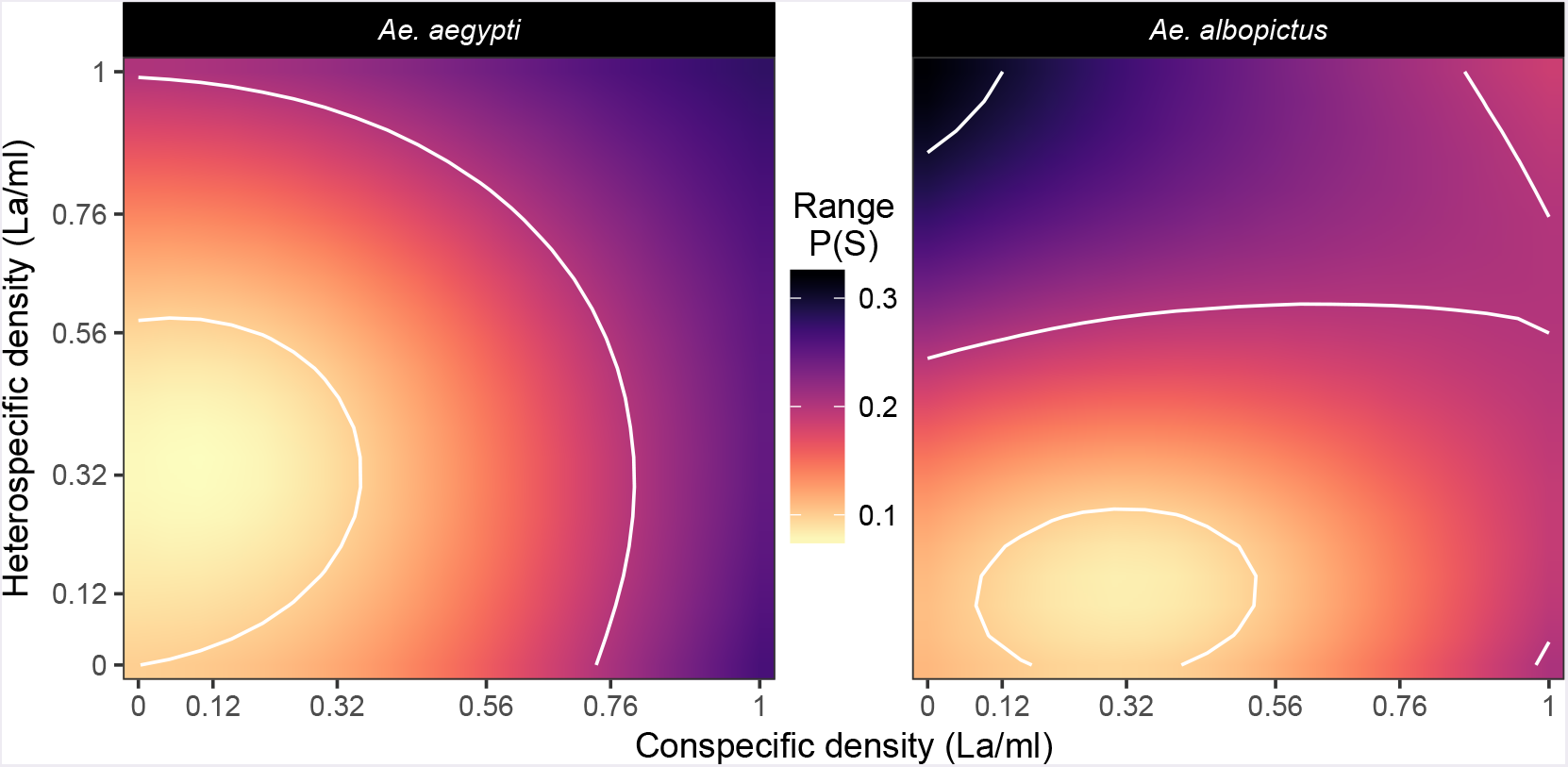
Plot showing the distribution of uncertainty (as a range in survival probability) across the predicted response surfaces in Figure 3.

**Figure. 11:**
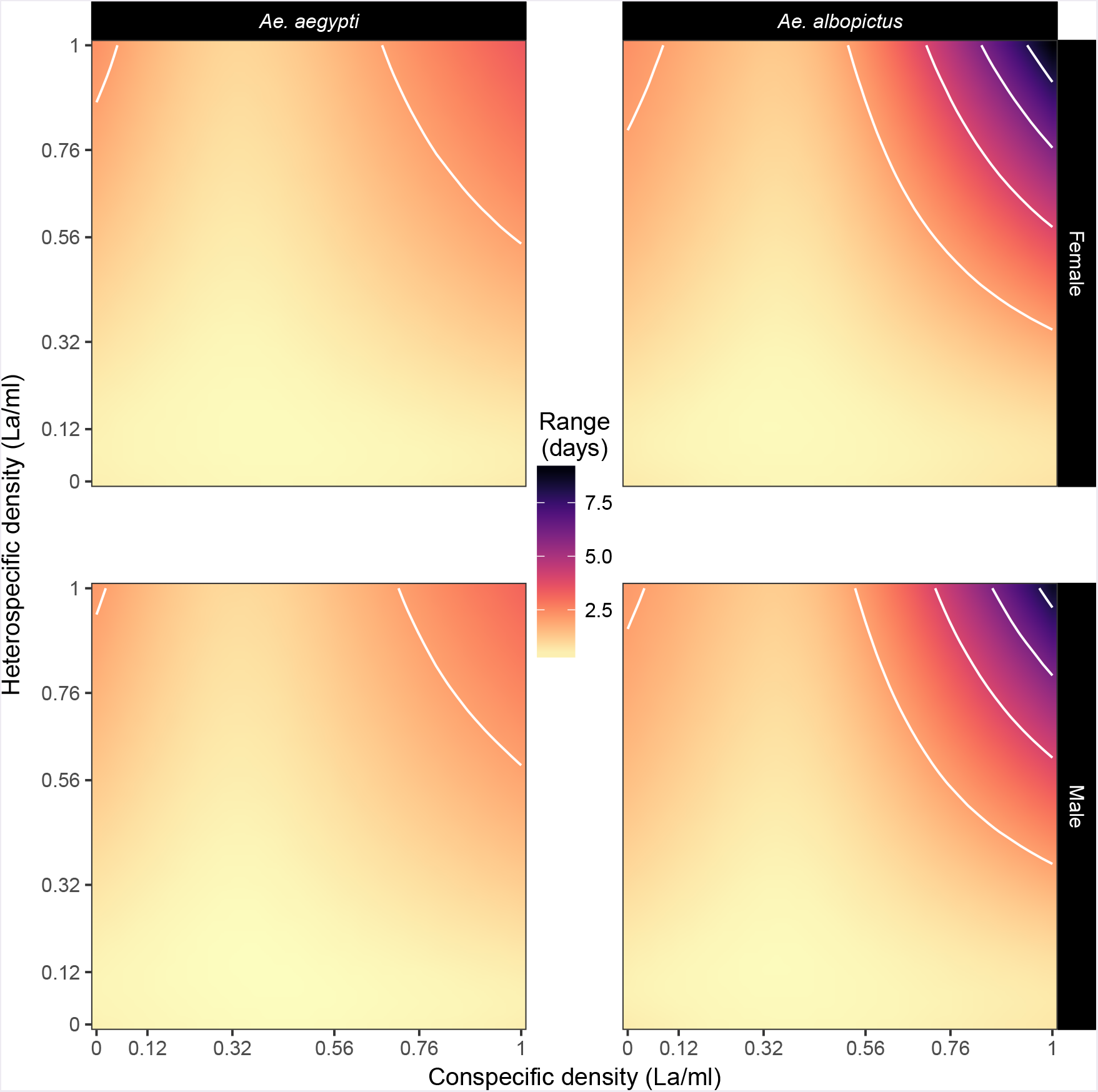
Plot showing the distribution of uncertainty (as a range in days) across the predicted response surfaces in Figure 5.

